# *A*2*G*^2^: A Python wrapper to perform very large alignments in semi-conserved regions

**DOI:** 10.1101/2020.05.21.109009

**Authors:** Jose Sergio Hleap, Melania E. Cristescu, Dirk Steinke

## Abstract

**Summary:** Amplicons to Global Gene (*A*2*G*^2^) is a Python wrapper that uses MAFFT and an “Amplicon to Gene” strategy to align very large numbers of sequences while improving alignment accuracy. It is specially developed to deal with conserved genes, where traditional aligners introduce a significant amount of gaps. *A*2*G*^2^ leverages the *add sequences* option of MAFFT to align the sequences to a global reference gene and a local reference region. Both of these references can be consensus sequences of trusted sources. Efficient parallelization of these tasks allows *A*2*G*^2^ to align a very large number of sequences (> 500K) in a reasonable amount of time. *A*2*G*^2^ can be imported in Python for easier integration with other software, or can be run via command line.

**Availability:** *A*2*G*^2^ is implemented in Python 3 (3.6) and depends on MAFFT availability. Other package requirements can be found in the requirements.txt file at https://github.com/jshleap/A2G. *A*2*G*^2^ is also available via PyPi (https://pypi.org/project/A2G). It is licensed under the LGPLv3.

**Supplementary information:** Supplementary material is available at github as jupyter notebook.

## 1 Introduction

High throughput sequencing technologies resulted in an unprecedented surge of genomic data. This data explosion challenges many existing analysis pipelines that rely on global multiple sequence alignments (MSA) (Nishimura *et al*., 2016; Garriga *et al*., 2019). Although alignment-free methods sequence comparisons exist (Ren *et al*., 2018), plenty of software still require a global alignment of all sequences (i.e. phylogenetic software, some taxonomic assignment tools). Unfortunately, the most robust sequence aligners to date (i.e. MAFFT, MUSCLE, etc; Thompson *et al*., 2011) cannot align more than a few thousand sequences reliably and within a reasonable amount of time. In fact, it has been shown that alignment quality decays with an increasing number of sequences (Sievers et *al*., 2011). Newer software such as PASTA (Mirarab *et al*., 2015) and the regressive alignment algorithm (RAA) (Garriga *et al*., 2019) leverage traditional MSA software capabilities to create alignments for hundreds of thousands to millions of sequences. However, these strategies also suffer from the same weaknesses (Figure S2). Empirical evidence suggests that for more conserved regions, traditional MSA programs produce too many gaps in the alignment, introducing noise to downstream analysis (Figure S2; Golubchik et al., 2007). A prominent example for a conserved region is the DNA barcode Cytochrome Oxidase subunit I (COI) which has become the main marker for metazoan species identification. Repositories such as the Barcode Of Life Data-system (BOLD) (Ratnasingham and Hebert, 2007) hold millions of COI sequences creating the need for an alignment strategy that can handle thousands of sequences from conserved regions without adding erroneous gaps.

Another challenge for modern alignment algorithms is the increasing amount of short sequence reads (50-300bp) produced by most high-throughput sequencing platforms (Illumina and Ion Torrent). Often those sequences are the product of prior PCR amplification (amplicons) and need to be aligned to specific genomic regions, especially when mined from databases. Such short reads could force an alignment algorithm to offset sequences to a shorter set causing a misalignment.

As semi-conserved regions are often used in large scale phylogenetic analyses or for taxonomic assignments, there is a need for a tool that can both solve the gappy alignment problem for very large numbers of sequences, and identify and remove poorly aligned sequences utilizing commonly available computational resources. Here we present the python wrapper *A*2*G*^2^ as an algorithm to overcome these issues.

## 2 Implementation

### 2.1 Anchored alignment

The *A*2*G*^2^ algorithm was developed on the assumption that when aligning a set of sequences of a known genomic region (global target), it is necessary to provide reference for both local targets (amplicons) and global targets. A global target sequence, which can be a single sequence from a gene or region of interest, or a consensus sequence of said region, is aligned to a local target. The latter can also be a single sequence or a consensus sequence representing the target region. This local reference will serve as an anchor for the remaining sequences to be aligned to the global target (Figure 1). This strategy allows offset sequences to align with a different region of the global target, preventing them from adding noise on the alignment in the desired region.

**Figure 1:**
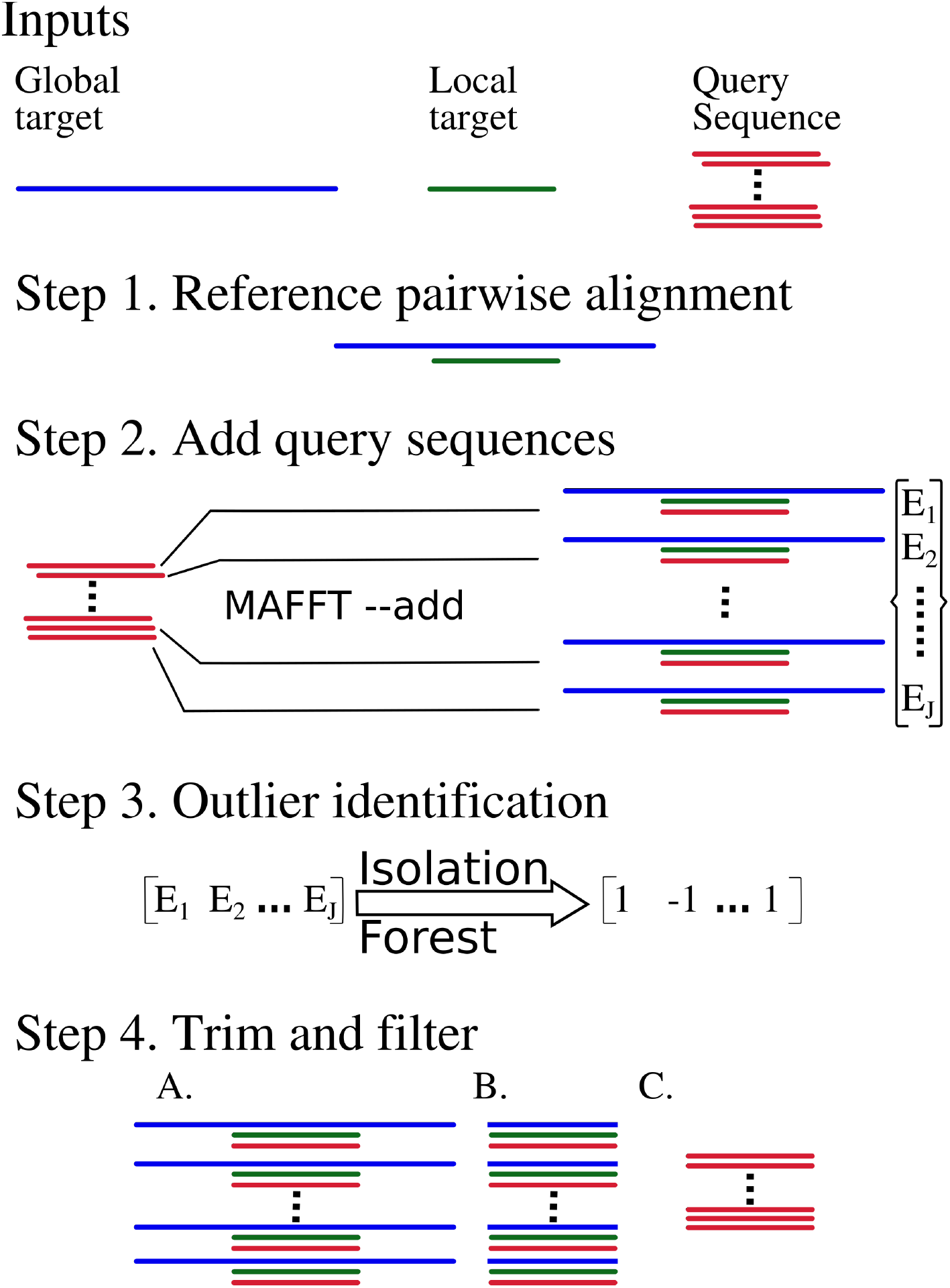
Schematics of the steps involved in *A*2*G*^2^ algorithm. *E_j_* is the median entropy from alignment *j*.

In the first step, the algorithm creates a pairwise alignment between the local and the global reference (Figure 1). Next, one query sequence at a time is added using a standard progressive alignment method implemented in the `MAFFT` option `--add` (Katoh and Frith, 2012). This step is run in parallel through the usage of multiple threads and CPUs or through a message passing interface in high performance computing environments. For each iteration of the alignment process, only the region aligned to the local reference is kept, the alignment is trimmed and aligned query sequences are stored.

### 2.2 Outlier detection and removal

To detect poorly aligned sequences (outliers), *A*2*G*^2^ uses the median Shannon entropy (*S*) across columns in the reference alignment (the alignment of the targets and *one* query sequence). We first estimate the frequencies of each base (in the case of nucleotides) or residue (in the case of proteins) in each column of each individual anchored alignment. Let *C*(*x*), be the counts of a base/residue *x*, in column *i*, and 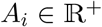, be the number of items in column *i*, then an entry in the array *p* will be defined as

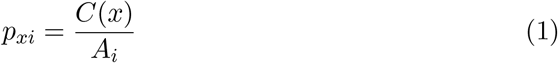

*p_xi_* then being the probability of base/residue *x* in the alignment column *i*. Now, for each column *i* we compute the Shannon entropy (*Si*) as:

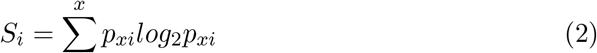

We then compute the median of all *i* columns present in the reference alignment of query sequence *j* to the global and local targets, as

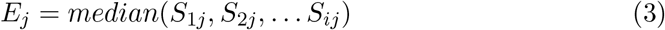

We apply equation 3 to all *j* query sequences, thereby obtaining the array *ϵ* = {*E*_1_, *E*_2_ … *E_j_*} which becomes the input for the outlier detection method known as isolation forest (Figure 1). Isolation forest is classified as an unsupervised learning algorithm for anomaly detection. This strategy is based on explicitly isolated anomalies instead of profiling normal points (Liu et al., 2008; Cheng et al., 2019). Isolation forests are based on the premise that anomalies or outliers are more easily isolated through random partition of a sample. Such a recursive partition can be represented as a tree structure known as isolation tree, hence the name isolation forest. *A*2*G*^2^ uses the implementation of isolation forest available in Scikit-learn (Pedregosa et al., 2011). Once the isolation forest method has identified entropic outliers, *A*2*G*^2^ will remove those query sequences from the alignment on request.

## Acknowledgements

The authors thank Jitka M. Krejci for comments on this manuscript. To Compute Canada for access to the Graham supercomputer for the scalability tests.

## References

Cheng, Z., Zou, C., and Dong, J. (2019). Outlier detection using isolation forest and local outlier factor. In Proceedings of the Conference on Research in Adaptive and Convergent Systems, pages 161–168.

Garriga, E., Di Tommaso, P., Magis, C., Erb, I., Mansouri, L., Baltzis, A., Laayouni, H., Kondrashov, F., Floden, E., and Notredame, C. (2019). Large multiple sequence alignments with a root-to-leaf regressive method. Nature biotechnology, 37(12), 1466–1470.

Golubchik, T., Wise, M. J., Easteal, S., and Jermiin, L. S. (2007). Mind the Gaps: Evidence of Bias in Estimates of Multiple Sequence Alignments. Molecular Biology and Evolution, 24(11), 2433–2442.

Katoh, K. and Frith, M. C. (2012). Adding unaligned sequences into an existing alignment using mafft and last. Bioinformatics, 28(23), 3144–3146.

Liu, F. T., Ting, K. M., and Zhou, Z.-H. (2008). Isolation forest. In 2008 Eighth IEEE International Conference on Data Mining, pages 413–422. IEEE.

Mirarab, S., Nguyen, N., Guo, S., Wang, L.-S., Kim, J., and Warnow, T. (2015). Pasta: ultra-large multiple sequence alignment for nucleotide and amino-acid sequences. Journal of Computational Biology, 22(5), 377–386.

Nishimura, Y., Amagasa, T., Inagaki, Y., Hashimoto, T., and Kitagawa, H. (2016). A system for phylogenetic analyses over alignments of next generation sequence data. In 2016 10th International Conference on Complex, Intelligent, and Software Intensive Systems (CISIS), pages 230–237. IEEE.

Pedregosa, F., Varoquaux, G., Gramfort, A., Michel, V., Thirion, B., Grisel, O., Blondel, M., Prettenhofer, P., Weiss, R., Dubourg, V., et al. (2011). Scikit-learn: Machine learning in python. Journal of machine learning research, 12(Oct), 2825–2830.

Ratnasingham, S. and Hebert, P. D. (2007). Bold: The barcode of life data system (http://www.barcodinglife.org). Molecular ecology notes, 7(3), 355–364.

Ren, J., Bai, X., Lu, Y. Y., Tang, K., Wang, Y., Reinert, G., and Sun, F. (2018). Alignment-free sequence analysis and applications. Annual Review of Biomedical Data Science, 1, 93–114.

Sievers, F., Wilm, A., Dineen, D., Gibson, T. J., Karplus, K., Li, W., Lopez, R., McWilliam, H., Remmert, M., Söding, J., et al. (2011). Fast, scalable generation of high-quality protein multiple sequence alignments using clustal omega. Molecular systems biology, 7(1).

Thompson, J. D., Linard, B., Lecompte, O., and Poch, O. (2011). A comprehensive benchmark study of multiple sequence alignment methods: current challenges and future perspectives. PloS one, 6(3).

